# A modular, high-bandwidth, bidirectional implantable device for neural interrogation

**DOI:** 10.64898/2026.02.23.707488

**Authors:** Radu Darie, Samuel R. Parker, Jonathan S. Calvert, Ekta Tiwari, Naser Abdelrahman, Sohail Syed, Elias Shaaya, Jared S. Fridley, Mark W. Merlo, Ian Halpern, David A. Borton

## Abstract

Modern neuroelectronic interfaces have shown great potential to diagnose conditions, address neurological dysfunction, and advance neuroscientific knowledge. However, neural interface systems today require tethered connections that restrict mobility, prevent testing across ecological contexts, and inhibit clinical translation to at-home use. Fully implantable commercial systems have previously been developed, but exhibit significant constraints, including a bulky design, limited modularity, low bandwidth, or unidirectional communication (e.g. deep brain stimulation systems, DBS; spinal cord stimulation systems, SCS). Here, we have developed the Modular Bionic Interface (MBI), a system composed of a fully implantable device and a worn unit for high-bandwidth, bidirectional interfacing with the nervous system. The MBI can record high fidelity electrophysiological signals and deliver spatiotemporally modulated electrical stimulation for clinical and research purposes through flexible interaction with third party implantable devices. We performed benchtop evaluation to validate the recording and stimulation capabilities of the MBI across a diverse range of inputs and outputs. We then evaluated the MBI system *in vivo* through chronic implantation within a sheep, where results were stable for the length of evaluation, over three months. While connected to an actively powered, third-party high-resolution spinal cord stimulation electrode array, the MBI system was able to deliver stimulation to evoke lower extremity motor responses and record spinal compound action potentials evoked by peripheral nerve and spinal stimulation. Through rigorous evaluation, we demonstrate a fully implantable system with a small footprint capable of high-resolution, bi-directional communication with the nervous system via modular connections to third-party devices. We expect that modular devices will further our ability to treat complex neurological disease and injury.

## Introduction

Neural interfaces have shown remarkable progress towards enabling the treatment of neurological disease, disorder, and injury [1]. Significant investment by governments, institutions, and researchers has led to an unprecedented array of devices and technologies to interact with the central nervous system (CNS) and peripheral nervous system (PNS) [2]. Modern neural interfaces range from 1000+ channel probes used in rodents, non-human primates, and humans [3]–[7], to molecular tools able to perturb and monitor cell-specific activity across space and time [8], transforming our ability to perform neuroscience research to understand and treat neurological disease and injury. A device that can support the advanced needs of future research must strike a delicate balance between mutually antagonistic constraints. Before describing one implementation of such a device, we will begin with a brief survey of the design space for neural interfaces and the many trade-offs it involves.

First, there is the question of where in (or on) the body to deploy probes in a way that maximizes therapeutic benefits while minimizing risk. Devices with invasive surgical placement involve higher risk of infection as well as treatment costs, but are also able to detect more precise measurements of disease state (e.g. intracortical neural action and local field potentials versus signals that can be obtained non-invasively, such as electroencephalography) and cause changes to neural activity on a finer spatiotemporal scale [17]. In terms of patient perspective, the potential benefits of implanted neural interfaces, which could be less obtrusive and provide users with more independence across activities of daily living than their wearable counterparts, may well outweigh the downside of requiring surgery [18]–[21].

Then, there is a related trade-off between device architectures that involve a percutaneous connection to external hardware and fully implantable devices. To date, systems developed for electrical neural interrogation (e.g. the Ripple Neuromed Summit or Blackrock Neurotech NeuroPort) have required percutaneous connections to extract high-fidelity information from, or provide high-resolution modulation of, neural systems and circuits. However, a percutaneous connection hinders experimentation across ecologically relevant contexts, such as a neuroprosthesis user’s home [9]. A percutaneous connection additionally introduces a persistent risk of infection and could severely impact a user’s mobility, depending on the specific use case. On the other hand, fully-implantable systems for neural interrogation are being developed [15]–[21], but have limited ability to provide interactions with the nervous system that are at once high-bandwidth, high-resolution and bidirectional. For fully implantable systems, wireless power and data transfer (WPDT) represents a significant bottleneck for performance. Data rate and power consumption are directly related, in that power requirements scale proportionally with channel count, analog-to-digital converter (ADC) resolution, and sampling rate. Additionally, power consumption imposes limits on device battery life and safety due to tissue heating, the latter being an important concern given current guidelines that require that implantable medical devices cause no more than a 2° C increase in temperature of the surrounding tissue [22]. As a result, research on the efficient delivery of power to devices located deep within the body has made great strides in recent years. While an in-depth exploration of WPDT is beyond the scope of the current publication, we refer the reader to excellent reviews on these topics, present in the literature: [23]–[25].

Finally, modularity and the ability for a flexible deployment are often overlooked but crucial elements of the ideal neural interface. For the reasons we have outlined thus far, neural engineering represents a massive systems integration challenge. Components of a device that could, in principle, be multipurpose, such as the electronics that support signal recording and processing, electrical stimulation, power management, or data telemetry are often developed at great cost as part of a single-target product that cannot readily be used to treat other indications. For example, devices with built-in electrodes for epicortical recording, such as the WIMAGINE system [26] have high spatiotemporal recording capabilities, but cannot be used for spinal cord recordings simply because the electrodes are not anatomically compatible. There is a need for a system that can be used agnostically of the target tissue, particularly for the field of psychiatric neuromodulation, where there is growing evidence that treatments must target widely distributed brain networks [27], or for motor rehabilitative spinal cord stimulation that needs to span several myotomes [28]. In summary, to develop a device that can enable the next generation of advances in neurotechnology we must find an equitable compromise between many competing requirements. Towards that goal, here, we introduce the Modular Bionic Interface (MBI), a system for high-bandwidth bidirectional interfacing with the nervous system (fig. 1, 2) composed of a fully implantable device (fig. 2a) and a worn unit for power and data telemetry (fig. 2b). The device is capable of recording high fidelity electrophysiological signals from micro- and macro-electrodes implanted in the central or peripheral nervous system. Additionally, the MBI can deliver spatially and temporally modulated electrical stimulation for a variety of clinical and research purposes. The modular design of the MBI enables flexible configurability with third party implantable devices. For example, we demonstrate the MBI connected to an actively multiplexed spinal cord stimulation electrode array for high resolution, multichannel, spinal electrophysiology and neuromodulation (6464, Micro-Leads Medical) [29], [30]. Herein, we describe the overall design and architecture of the MBI, demonstrate its performance on the benchtop, and report the functionality of the MBI when chronically implanted in a large animal model (Polypay sheep, *Ovis aries*) over a period of three months.

**Figure 1.**
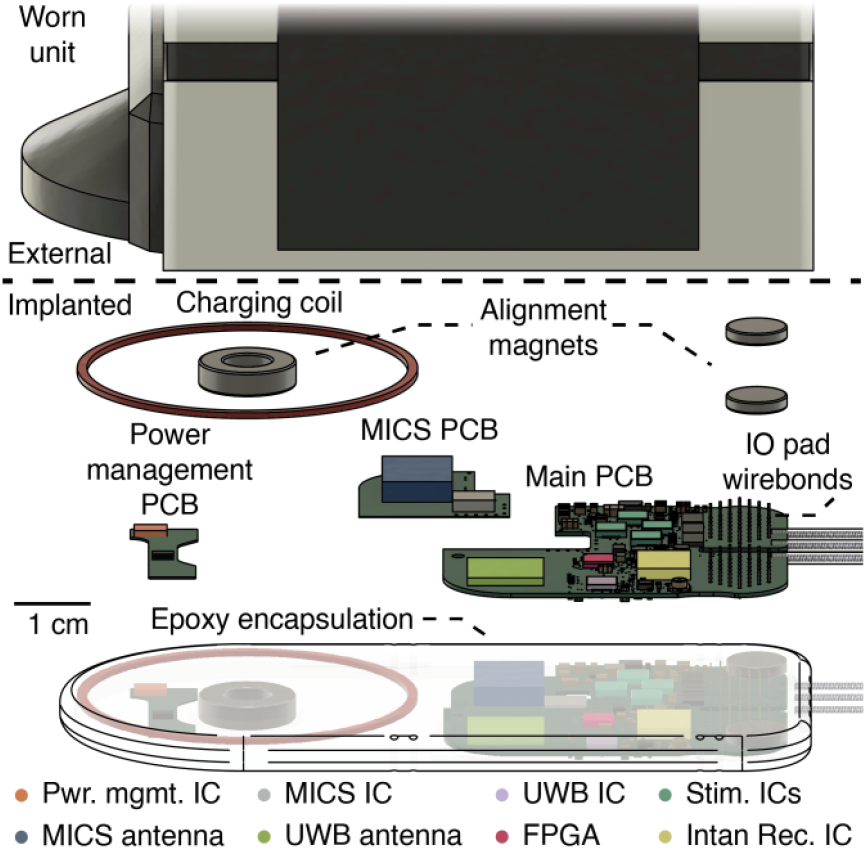
CAD illustration of the MBI system components (worn unit partially cropped out). The main subcomponents of the MBI implant are vertically offset and labelled. Important ICs on the implanted unit are labelled and false colored to match the outlines in figure 2. Abbreviations: CAD – Computer Aided Design, FPGA - field programmable gate array, IC - integrated circuit, MICS - medical implant communication system, UWB - ultra wide band, PCB – Printed Circuit Board

**Figure 2.**
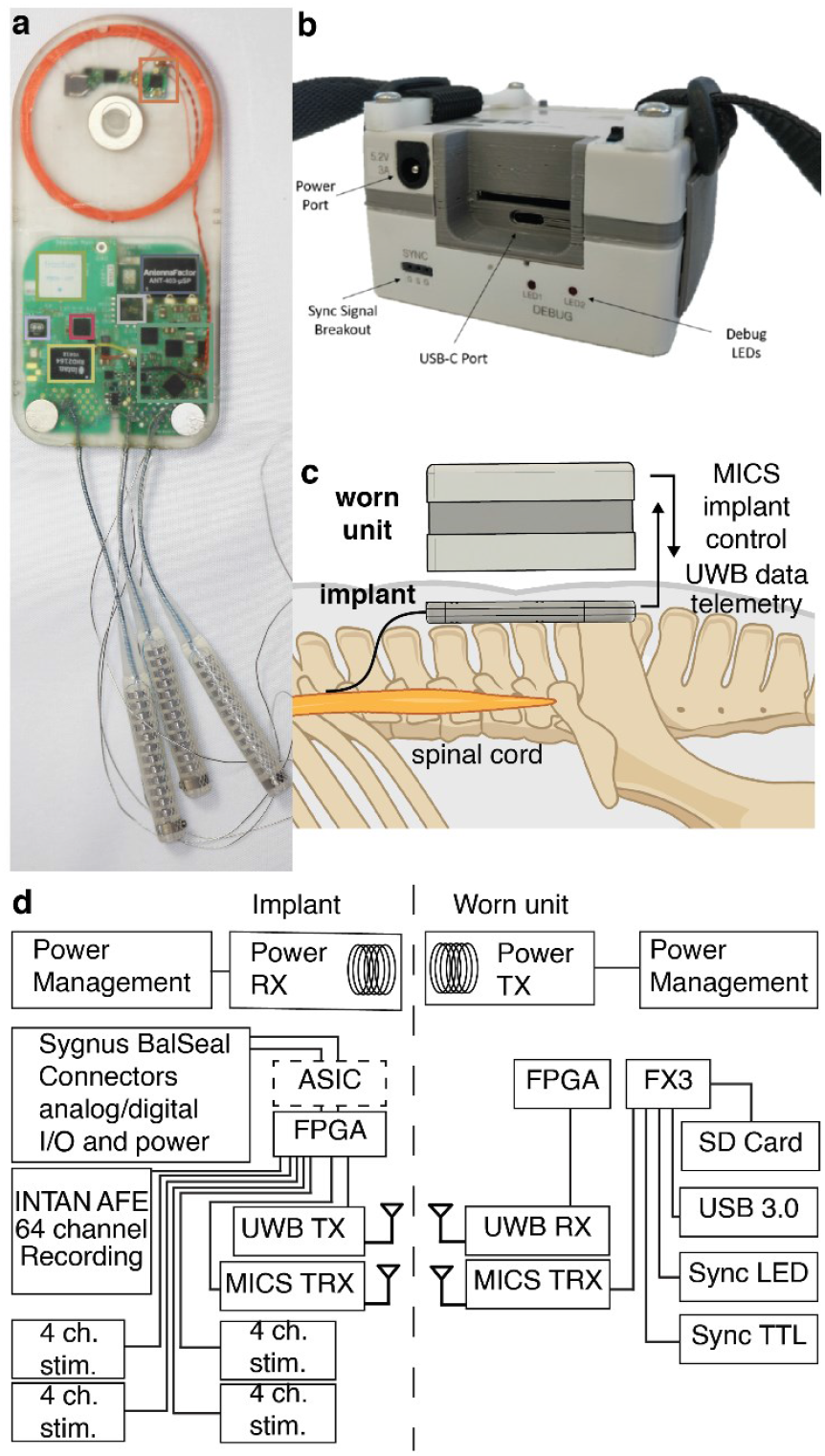
MBI system illustration. (a) Photograph of the MBI implant. (b) Photograph of the worn unit. (c) Cartoon schematic depicting one possible use of the MBI to interface with a device implanted in the spinal cord. (d) System diagram of the MBI implant and worn unit. The implementation described in this text features an optional control ASIC (dashed outline). Abbreviations: AFE - analog front end, ASIC – application specific integrated circuit, MICS - medical implant communication system, UWB - ultra wide band, FPGA - field programmable gate array, TX - transmit, RX - receive, TRX - transmit and receive.

## Methods

### System architecture

As a platform, the functionality of the MBI is split between the implanted unit itself (fig. 2a) and an external wearable unit (fig. 2b), ensuring that the implant remains thin, with a small footprint, for easy surgical implantation and reduced risk of infection.

The worn unit magnetically aligns to the implant and provides power wirelessly, through an inductive link, eliminating the need for an implanted battery. Furthermore, the worn unit handles power management, bi-directional wireless communication, and communication with a host computer via USB 3.0. All digital operations on the worn unit are split between a field-programmable gate array (FPGA) running custom firmware and a high-speed USB peripheral controller (EZ-USB™ FX3, Infineon, Munich, Germany) (fig. 3d). Writing data to a removable SD card was implemented for future long-term experiments but was not used in the current demonstration. Additionally, the worn unit features a TTL output and LED indicator light that flash at a rate of 1 Hz, for synchronization with other electrophysiology systems. The host computer controls recording and stimulation via a high-level C++ API that can be extended to suit the needs of individual experiments.

**Figure 3.**
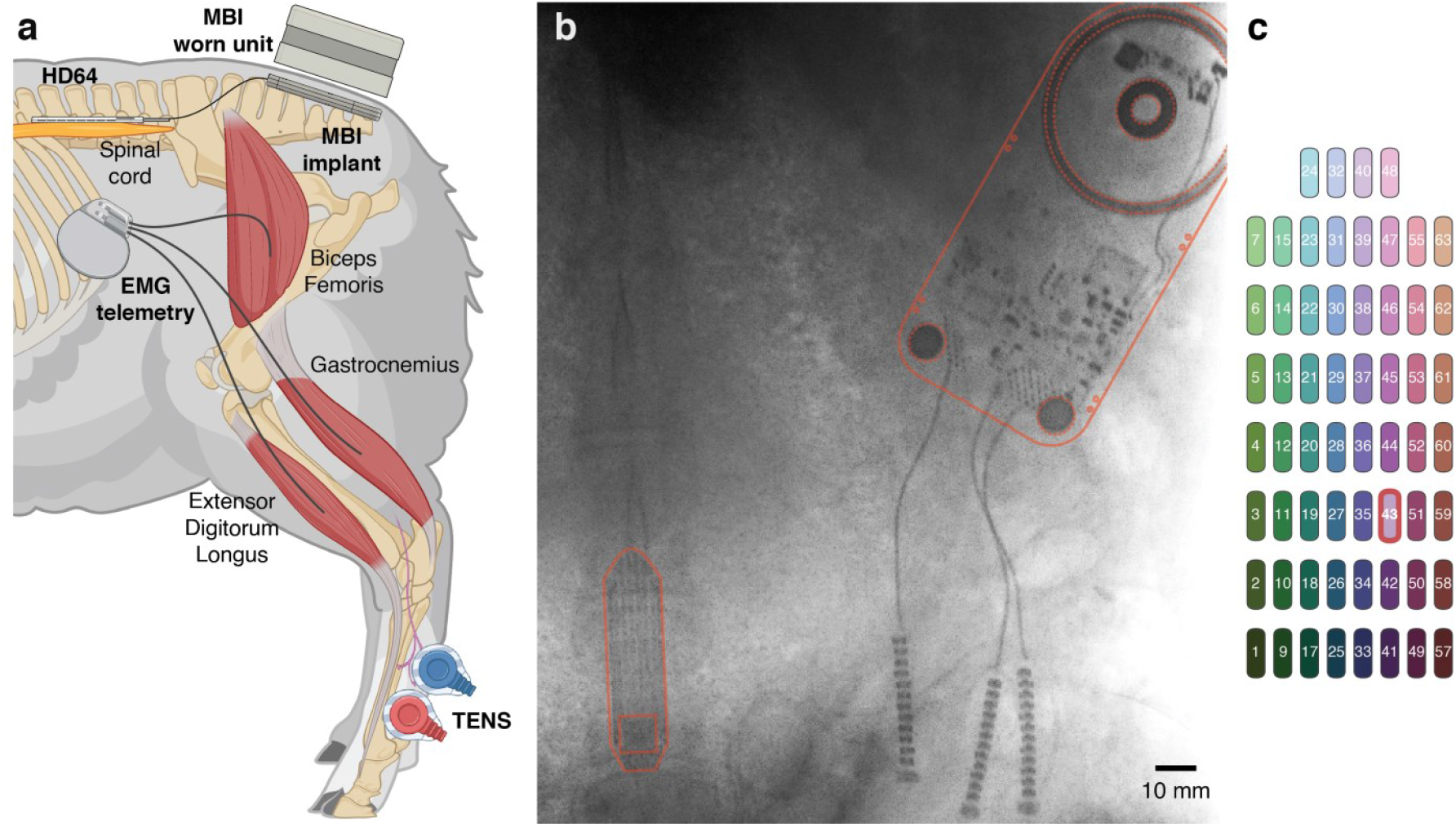
In vivo experimental setup. (a) Cartoon illustration of the animal model used for in-vivo validation of the MBI. The MBI is implanted subcutaneously and connected to an epidural electrode array for spinal cord stimulation and recording. The worn unit rests on the animal’s skin and magnetically aligns to the MBI. Separate systems are used for EMG recording (implanted) and TENS (external), respectively. (b) X-Ray image of the implanted MBI unit and epidural electrode array, outlined in red for ease of visualization. (c) Topographic representation of all recording channels on the spinal electrode array. For each experimental session, we selected 12 of these electrodes as recording channels, as indicated in the legends of figures 6 and 7. Electrode 43, used to provide stimulation (figure 7), is indicated by a red outline.

The implanted unit houses the circuitry for recording, stimulation, signal processing, wireless power, and data transfer (fig. 2d). All data operations on the MBI implant are performed by an FPGA (ICE40LP8K-CM81, Lattice Semiconductor, Hillsboro, Oregon, USA) that digitally controls recording, stimulation and data telemetry. The implanted unit is designed to maximize compatibility with passive or active implanted arrays of electrodes, or, in general, other active implanted devices. For clarity, we will refer to these as “distal” implants and to the MBI implanted unit as the “proximal” implant. Two fundamental features of the MBI’s design enable its use with different distal implants over a wide variety of experimental scenarios: 1) the ability to reprogram the FPGA; and 2) the presence of an array of 102 input-output connection points (IO pads). 64 of these IO pads can be used for simultaneous monopolar voltage recording, 16 can be used to source and/or sink stimulation current, while 22 are used for grounding, signal reference, power delivery or digital communication with the distal implant(s). The system can be adapted to specific scenarios by configuring, as needed, the IO pad connection routing, the FPGA firmware, or, in advanced cases, by adding custom components to the proximal implant circuitry. In the relatively simple scenario of using a passive electrode array, connections can be made by directly welding each electrode to a recording or stimulation IO pad. The same routing could be achieved by interposing an implanted connector [31]. In the implementation described in this paper, a subset of the IO pads are welded to the ends of 3 cylindrical Sygnus connectors, with 24 contacts each (Bal Seal, Foothill Ranch, CA). One of the Sygnus connectors is wired to 24 recording inputs, enabling passive recording from pre-existing electrode designs used in clinical pacemakers and neuromodulation devices. The remaining two connectors are configured for a mixed digital and analog interface with the Micro-Leads HD64, an actively multiplexed spinal electrode array that offers access to 60 electrode contacts over just 24 signal wires [30]. The HD64 requires an application specific integrated circuit (ASIC) to configure and power the on-board multiplexer – the HD64 ASIC was added to the proximal implant PCB for the implementation described in this document. The control ASIC receives instructions from the FPGA and powers and configures the HD64 via digital IO pads to route the electrical connection between each spinal site and its requested recording/stimulation pads.

The MBI uses an Intan RHD2164 64-channel electrophysiology amplifier chip to acquire and digitize electrical signals from neural tissue. As a result of the low input-referred noise (3.12 μV_rms_ in CMOS signaling mode; see pg. 12 of [36] for details) and field-programmable bandwidth, these amplifiers are suitable for a wide range of neurophysiological recordings, with targets including single neuron action potentials, local field potentials, and evoked compound action potentials. The same MBI device can also be used to record other biopotentials, such as electromyography (EMG) or electrocardiogram (ECG). Being fully implanted, the MBI is isolated from environmental sources of electrical interference, resulting in high signal-to-noise compared to using an external recording system to acquire the same signals. An ultra-wideband (UWB) communication protocol enables the MBI to wirelessly transmit multiple channels of high-resolution raw data, and to resolve neural events on a sub-millisecond scale - e.g. putative single neuron action potentials (fig. 4c), or stimulation evoked compound action potentials (fig. 6, 7). UWB transmission is handled by a pair of custom ASICs developed in-house (Modular Bionics, CA) to transmit (on implant) and receive (on worn unit) data. The UWB ASICs were fabricated using a 65 nm liquid phase epitaxy complementary metal-oxide-semiconductor process (LPE CMOS). The telemetry system operates in the 3.1 GHz to 5.1 GHz band with a center frequency of 4.056 GHz. It implements a direct-sequence code division multiple access (DS-CDMA) protocol with a chipping rate of 1.352 GHz. The transmitter consumes 26 mW and has a raw and effective energy per bit of 250 pJ/bit and 269 pJ/bit, respectively. The communication protocol utilizes a packet structure with a 32-bit header CRC and 32-bit payload CRC for error detection and a redundant header for error correction. The telemetry system and protocol have been tested at speeds as high as 104 Mbits/sec (96.5 Mbits/sec effective raw data rate – discounting packet overhead) for higher channel count and/or temporal resolution than used in the *in vivo* implementation reported here. This specific implementation samples from 64 channels at a rate of 37 kHz, for a combined effective data rate of ∼38 Mbits/sec. However, for all experiments, we used the remaining bandwidth to transmit a dummy signal, consisting of all zeros, to ensure that data transmission remains reliable at the full 96.5 Mbits/sec rate.

**Figure 4.**
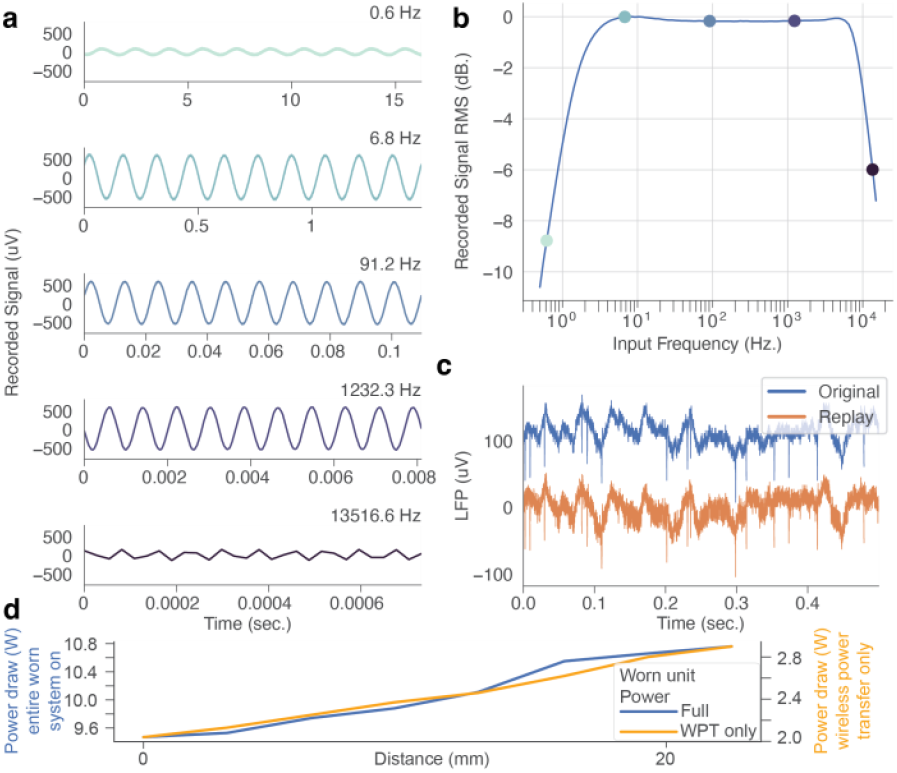
Benchtop evaluation of the MBI demonstrates stable recording performance. (a) Select windows of data from the frequency sweep recording, showing 10 cycles of the sinusoidal inputs at the indicated frequencies. (b) Bode plot illustrating the recording frequency response. Colored markers correspond to the frequencies illustrated in panel a. (c) Previously recorded intracortical LFP and the corresponding MBI recording. The traces are vertically offset for ease of visualization. (d) Worn unit power draw as a function of distance to the implant, with subsystems disabled (light orange trace, right axes), or fully enabled (dark blue trace, left axes), respectively. WPT – wireless power transfer.

The MBI uses a set of 4 quad-channel programmable current sink/source chips (CSI021, Cirtec Medical, CT) for up to 16 channels of simultaneous and independent electrical stimulation. These chips offer several advantages for neuromodulation applications. Stimulation parameters can be programmed independently on each channel, offering precise control over stimulation pulse amplitude (12 μA resolution) and timing (80 μs resolution for the inter-pulse period, and 10 μs resolution for pulse width). Stimulation pulses are automatically charge balanced, to protect from potential tissue damage associated with long-term charge accumulation. The combination of high parameter resolution and high compliance voltage range (−7.9 V to 1.7 V) covers a range of stimulation parameters commonly used for a variety of neural targets ranging from intracortical microstimulation [32] to epidural spinal cord stimulation [33]. Pulse parameters and timing are controlled by a host computer and relayed to the MBI via the worn unit over a medical implant communication system (MICS) band radio connection (ZL70323, Microchip Technology Inc., Chandler, Arizona, USA). Stimulation parameters are then programmed and delivered via a high-speed serial peripheral interface (1-10 MHz), enabling fast real-time control of stimulation.

### Benchtop characterization

We first performed a series of benchtop experiments to confirm that the individual components of the system, which comprises custom silicon, custom firmware, and several off-the-shelf components, operate within their specified parameters when assembled into a complete device. Each are discussed below.

### Worn unit power consumption vs displacement from implant

To ensure that the device’s operation was robust to changes in worn unit position, we performed benchtop testing of worn unit power consumption as a function of displacement from the implant. The worn unit was powered by a benchtop power supply (E36312A, Keysight Technologies) set to 5.2 V. Worn unit power consumption was measured with the power consumption monitor of this power supply. The displacement between the worn unit and implant was controlled using one or more 3.2 mm thick acrylic spacers. For a displacement of 0 mm no acrylic spacers were used, and the worn unit and implant were contacting each other. The coils alignment at each displacement was maintained by the alignment magnets in the worn unit and implant, and this alignment was confirmed with visual inspection. The power consumption was recorded for two conditions: (1) while powering just the worn unit wireless power system (to power the implant) and (2) with the worn unit fully powered (for wireless data retrieval and processing) – Fig. 4d.

### Benchtop evaluation of recording and stimulation capabilities of the MBI

Next, we sought to confirm the fidelity of recording and stimulation with the MBI, in a benchtop setting. To characterize the overall frequency response of Intan recording chip when implemented on the MBI, we used a NeuroDAC [34], an arbitrary waveform generator capable of accurately producing microvolt-scale outputs, to generate a sweep of sinusoidal signals that we recorded on the MBI (fig. 4a). We generated 100 pure sinusoidal input signals at a resolution of 192kHz with an amplitude of 500 μV_pk-pk_ and frequencies logarithmically spaced between 0.5 Hz and 15 kHz. The duration of each sinusoidal segment was set to be no shorter than 3 seconds and span at least 100 cycles of the oscillation. To obtain the frequency response, we calculated the RMS amplitude of the recorded signal, normalized to the maximum observed value in the passband and converted the result to decibels (fig. 4b).

We generated and recorded a copy of previously recorded intracortical local field potentials from somatosensory area 3a of a nonhuman primate (*Macaca mulatta*). The original time series was recorded from a chronically implanted microelectrode array (N-Form, Modular Bionics, CA) while the animal was awake and stationary in a standard primate chair. The raw data was originally sampled at 30 kHz using a Nano2 headstage (Ripple Neuro, Salt Lake City, UT), resampled to 192 kHz for NeuroDAC playback and recorded on the MBI at 36.932 kHz (Fig. 4c).

To capture the stimulation pulse waveform, we connected an RC circuit representing the tissue-electrode interface [35] across 1 (fig. 5a, 5b) or 2 (fig. 5c) MBI stimulation outputs and recorded the voltage across the circuit using a digital oscilloscope (Analog Discovery 2, Digilent, Pullman, WA). Stimulation pulses were delivered at a constant 1200 μA amplitude with pulse widths spanning 10 - 1380 μsec (fig. 5a), or a constant 100 μsec pulse width with amplitudes spanning 12 - 3060 μA (fig. 5b).

**Figure 5.**
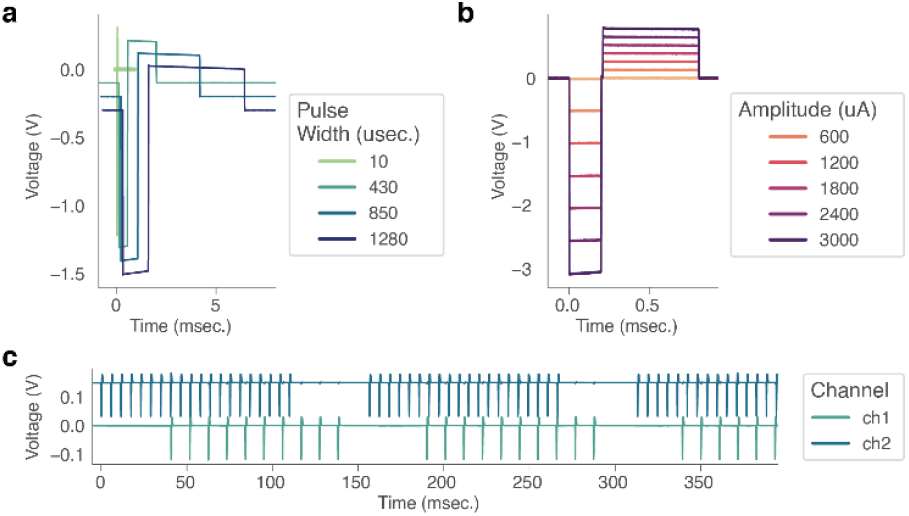
Benchtop evaluation of the MBI demonstrates stable stimulation performance. (a) Waveform corresponding to stimulation pulses delivered at 1200 μA and increasing pulse width. Traces are offset vertically and horizontally for ease of visualization. (b) Waveform corresponding to stimulation pulses delivered at 100 μsec pulse width and increasing amplitude. (c) Simultaneous recording of stimulation delivered at 1200 μA and a pulse width of 100 μsec, with pulse rates of 175 Hz (top trace, blue) and 90 Hz (bottom trace, green) respectively.

### Surgical procedure for animal experiments

#### Overview

All surgical and animal handling procedures were completed with approval from the Brown University Institutional Animal Care and Use Committee (IACUC), and in accordance with the National Institutes of Health Guidelines for Animal Research (Guide for the Care and Use of Laboratory Animals). One adult female Polypay sheep (*Ovies Aries*, approximately 90 kg and 3.5 years old at time of MBI implant) was used for this experiment. The animal was housed in a controlled environment on a 12-hour light/dark cycle with ad-libitum access to water and was fed twice daily. The animal was used for several experiments beyond the scope of this study, which required a staggered approach to the implantation of the epidural electrode array, MBI implant and EMG telemetry unit. All surgical procedures followed the same general flow, as follows.

On the day of each surgery, the animal was initially anesthetized with a mixture of ketamine, midazolam, and Xylazine, then intubated and transitioned to isoflurane (1-5%) inhalation. The skin over the surgical site was disinfected with a surgical scrub and the sheep was positioned prone on the operative table. The sheep was placed in a V-shaped foam block to maintain proper alignment of the spine, and a rectangular foam block was placed under the hip to alleviate pressure on the hind legs. When necessary, to enable intraoperative electrophysiological measurements, the animal was transitioned temporarily from isoflurane to intravenous anesthesia (Propofol 300 ug-600 ug/kg/min + Fentanyl 0.15-0.225 ug/kg/min).

At the end of each surgery, incisions into fascia and superficial layers were closed with resorbable Vicryl suture. Skin incisions were closed with resorbable Monocryl suture and dressed with Dermabond and a sterile dressing. Finally, the sheep was transferred to a sling (Panepinto, Fort Collins, CO) and monitored as it emerged from anesthesia. Once the animal fully recovered, they were returned to their pen.

#### Epidural electrode array implantation

The epidural electrode array (HD64, Micro-Leads Medical, Somerville MA) was implanted first and used for other experiments with an external electrophysiology system through percutaneous connections. The L5-L7 spinous processes were identified using anatomic landmarks, and a linear skin incision was made sharply using a scalpel. Once the L5-L7 spinous processes, laminae, and facet joints were exposed bilaterally, a laminectomy with medial facetectomy was performed. The electrode array was gently placed over the exposed thecal sac. The electrode leads were then secured to the paraspinal muscles, and a stress relief loop was made. Each of the two leads was connected to a standard extension lead (Nuvectra, Plano, TX), which received a second strain relief loop. The extensions were then tunneled laterally and externalized through a small circular incision in the skin. Custom-made ground wires, constructed from Cooner AS636 wire (Cooner Wire Company, Chatsworth, CA), were placed epidurally and in the paraspinal muscles that were previously dissected off the lamina, tunneled laterally and externalized through the same opening as the extension leads. Adequate strain relief was ensured for the ground wires as well. At the end of the procedure, stimulation was delivered to confirm optimal array positioning by stimulating at multiple spinal locations and observing lower extremity motor activation.

#### EMG recording system implantation

Two PhysioTel L03 telemetry implants (DSI Systems, St. Paul, MN) were implanted bilaterally to obtain EMG recordings from the left and right Biceps Femoris, Gastrocnemius and Extensor Digitalis Longus muscles. The surgical preparation was similar to the other procedures, with the exception that the hind legs were allowed to drape over the sides of the surgical table, enabling access to the hind leg musculature. Additionally, the entirety of the hind legs was shorn and sterilized, creating a wide surgical field. Incisions were made roughly 10 cm anterior to the iliac crest and 40 cm laterally on each side. Subcutaneous pockets were developed for the placement of the implant main units. The location of target muscles was determined by palpation. The muscles were then exposed by sharp incision, and their identity was confirmed visually. EMG recording leads, extending from the implant main units, were tunneled subcutaneously to their target locations by blunt dissection. Each recording lead (two per muscle for bipolar recording) consists of a silicone insulated stainless steel wire. 5 mm lengths of wire were exposed and placed inside the muscle using a 16-gauge needle. First, the needle was tunneled through the muscle belly. Then, the wire was placed into the lumen of the needle such that, as the needle was retracted, the wire followed and embedded itself into the muscle belly. Once in place, the leads were secured to the muscle fascia using non-resorbable suture. Once all EMG leads were in place, all incisions were closed.

#### MBI implantation

An incision was made to expose the electrode and extension leads. A location was identified roughly 20 cm cranial and 20 cm lateral from the connectors, where a second incision was made. A subcutaneous pocket was made by blunt dissection and the MBI was placed within. The Bal Seal connectors were tunneled to the first incision. Here, the electrode leads were disconnected from the extensions and reconnected to the native Bal Seal connectors. The percutaneous extensions and ground wires were removed. The MBI ground and reference wires were secured to the nearby muscle fascia by non-resorbable suture. At the end of the procedure, stimulation was delivered through the MBI to confirm device operation, by observing lower extremity motor activation. Note that, while the MBI is capable of recording EMG, we did not make use of that functionality in this study, to eliminate the added surgical burden of tunnelling and placing a new set of EMG leads.

#### In vivo experimental procedures

At the beginning of sling-based experimental sessions, the fully conscious sheep was hoisted in a sling (Panepinto, Fort Collins, CO) until clearance between its hooves and the floor was observed, to ensure that its weight was fully supported by the sling. The worn unit was connected to a control PC via USB, then magnetically aligned and attached to the MBI. Electrode routing, data capture and stimulation control were controlled using software developed in-house at Modular Bionics. EMG data was streamed continuously throughout the experimental session at 500 Hz using the Ponemah software suite (DSI Systems, St. Paul, MN). The MBI synchronization wave was sampled at 1000 Hz using a wired connection between a DSI analog signal interface and the worn unit. Throughout the duration of the experiment, the animal was monitored for signs of discomfort or stress and was fed grain intermittently.

For spinal cord stimulation experiments, a subset of the 60 epidural electrodes was configured for recording or stimulation at the beginning of each experimental block. 300 msec trains of stimulation were delivered at 1-1.5 second intervals while the animal was stationary in the sling. All pulses had cathode-leading asymmetric biphasic charge balanced waveforms with a 1:4 ratio between the durations of the cathodic and anodic phases. While mono- and multi-polar cathode configurations were investigated, the return path for the current was always the subcutaneous ground wire. Other stimulation parameters were varied to investigate the effects of different combinations of stimulation amplitude, frequency, pulse-width and location on the recruitment of EMG and spinal extracellular compound action potentials (ECAPs).

For the peripheral nerve stimulation experiments, the bony anatomy of the hind fetlock was palpated, and a ring of wool just proximal to the fetlock was shorn, extending approximately 10 cm proximally.

The exposed skin was cleaned with isopropyl alcohol, which was allowed to air dry. 2” x 2” TENS (transcutaneous electrical nerve stimulation) patches (Balego, Minneapolis, MN) were placed on the medial and lateral aspects of the cannon bone, then connected to the external stimulator (Model 4100, A-M Systems Inc, Carlsborg, WA). The stimulation monitoring channel was connected to an analog input on the DSI signal interface for synchronization. Stimulation amplitude was applied at 15V, 20V, and 25V. In all cases, the stimulation pulses had cathode leading symmetric charge balanced waveforms with a width of 250 μs per phase. Single pulses were delivered with an interstimulus interval of 2.5 seconds. The above process was repeated for the left and right hindlimbs.

#### Data processing

All analyses of benchtop and *in vivo* data were performed in Python 3.8 using standard data processing packages (SciPy v1.10.0, NumPy v1.24.2, scikit-learn v1.2.1 and pandas v1.5.3).

#### Synchronization and signal processing

We coarsely synchronized the EMG and epidural spinal potential recordings based on the PC system timestamps associated with the respective raw data files. Next, we calculated the cross-correlation function between the two records of the synchronization wave, obtained from the DSI signal interface and original MBI data stream, respectively. We applied a temporal offset to the DSI system data at the lag of maximum cross correlation, to correct for inconsistencies between system clock timestamps and ensure millisecond-scale precision synchronization between the two data sources.

EMG data was high-pass filtered (8-pole Butterworth filter with a 0.5 Hz corner frequency) to remove movement artifacts, then z-scored. Spinal potential data was low-pass filtered (8-pole Butterworth filter with a 1 kHz corner frequency).

For the analysis of spinal cord stimulation evoked ECAPs (fig. 7), we employed a bipolar re-referencing scheme by taking the difference between signals recorded at vertically adjacent contacts.

## Results

### Benchtop characterization of the MBI

We performed a series of benchtop experiments whose goal was to validate that the recording and stimulation subsystems on the MBI were operating within the specifications of the RHD2000 series amplifier [36] and CSI series current source [37] as defined in their respective datasheets.

To characterize recording capability, we evaluated the system gain and spectral bandwidth (fig. 4). First, we observed that the frequency response of the recording was consistent with the bandpass filter options defined for the Intan RHD2164: a first order analog Butterworth high pass filter with a cutoff of 1 Hz, a third order analog Butterworth low-pass filter with a cutoff of 7.5 kHz, in combination with a digital high-pass filter with a cutoff of 1.44 Hz. No evidence of distortions in the signal chain was observed (Fig. 3a, b). To illustrate the ability of the MBI to capture low-amplitude, high-bandwidth signals, typical features of neurophysiological recordings, we recorded a copy of a previously obtained section of intracortical LFP that displayed both low frequency fluctuations (∼10 μV in amplitude) and evidence of single-neuron “spikes” (∼100 μV in amplitude, Fig. 3c). We observed that the MBI was able to faithfully reproduce these signals (fig. 4c). However, it proved difficult to meaningfully quantify the match between the original and the recorded signals, as the NeuroDAC itself cannot produce the low frequency content of the recording.

We tested the stimulation capabilities of the MBI by recording the stimulation pulse waveform across a range of pulse widths and amplitudes common for neural stimulation experiments (fig. 5). We saw agreement between programmed stimulation parameters and delivered stimulation. We swept pulse widths and amplitudes across a range of clinically relevant values: 10 microseconds to 1.28 millisecond pulse width (fig. 5a), 600 μA to 3 mA amplitude (fig. 5b). Note that the recorded voltage recordings exhibit slight decreases over the duration of each phase of the stimulation pulse, consistent with the charging of the capacitor used to model the electrode-tissue interface, indicating that the current injection followed the configured waveform (cathode leading biphasic pulses with a 1:4 cathodic-anodic pulse width ratio). Finally, we demonstrate one advantage of using multiple independent current sources: the ability to stimulate on different contacts simultaneously at different pulse rates (fig. 5c). Here, the device was configured to deliver 100 millisecond long pulse trains to two resistive loads at frequencies of 175 Hz and 90 Hz, respectively, which fall in and out of phase over time.

Benchtop testing of the wireless power system showed that the MBI worn unit can provide wireless power to the implant at displacement between 0 mm (touching) and 22.5 mm. When only the worn unit was powered, the power consumption started at 2.04 W at 0 mm displacement and increased with displacement to 2.9 W at 22.5 mm displacement. When the implanted unit received wireless power, the power consumption started at 9.47 W at 0 mm displacement and increased with displacement to 10.76 W at 22.5 mm displacement (fig. 4d).

### In-vivo characterization

We performed in-vivo characterization of the MBI while connected to a previously implanted epidural spinal cord electrode array (HD64) in a biologically relevant ovine model. The goal of these in-vivo experiments was to confirm the long-term functionality of the MBI once deployed in a living subject. Recording experiments highlighted the sensitivity necessary to capture biologically meaningful variations of the epidural spinal potential. Stimulation experiments highlighted the ability of the MBI to evoke neural activity, as evidenced by evoking muscle contractions (EMG) and spinal evoked compound action potentials (ECAPs).

### Evoked Potentials from Peripheral Nerve Stimulation

We used the MBI to record peripherally-evoked spinal ECAPs following transcutaneous electrical stimulation over the superficial peroneal and medial plantar nerves of the hindlimb fetlock (fig. 3a, [38]). Transcutaneous stimulation evoked complex, multiphasic evoked potentials (fig 6). The set of recording electrodes chosen for this experiment (see fig. 3 and the legend of fig. 6) span a wide medio-lateral (11.9 mm) and rostro-caudal (29.4 mm) range. Notably, while the responses are morphologically similar, there are marked differences depending on the location of the recording electrode, such as the amplitude of the initial negative deflection at around 10ms after the pulse. Simultaneous lower extremity EMG recording was performed to contextualize the spinal ECAPs with respect to the stimulation-evoked muscle activity. Additionally, the initial deflection in the ECAP occurs with a latency of approximately 10 msec, consistent with the expected propagation delay of action potentials from the nerve to the recording site (roughly 100 m/s). EMG responses, on the other hand, begin to occur with a latency of approximately 20 msec, suggesting that the muscle contractions are the result of a spinally mediated reflex arc.

**Figure 6.**
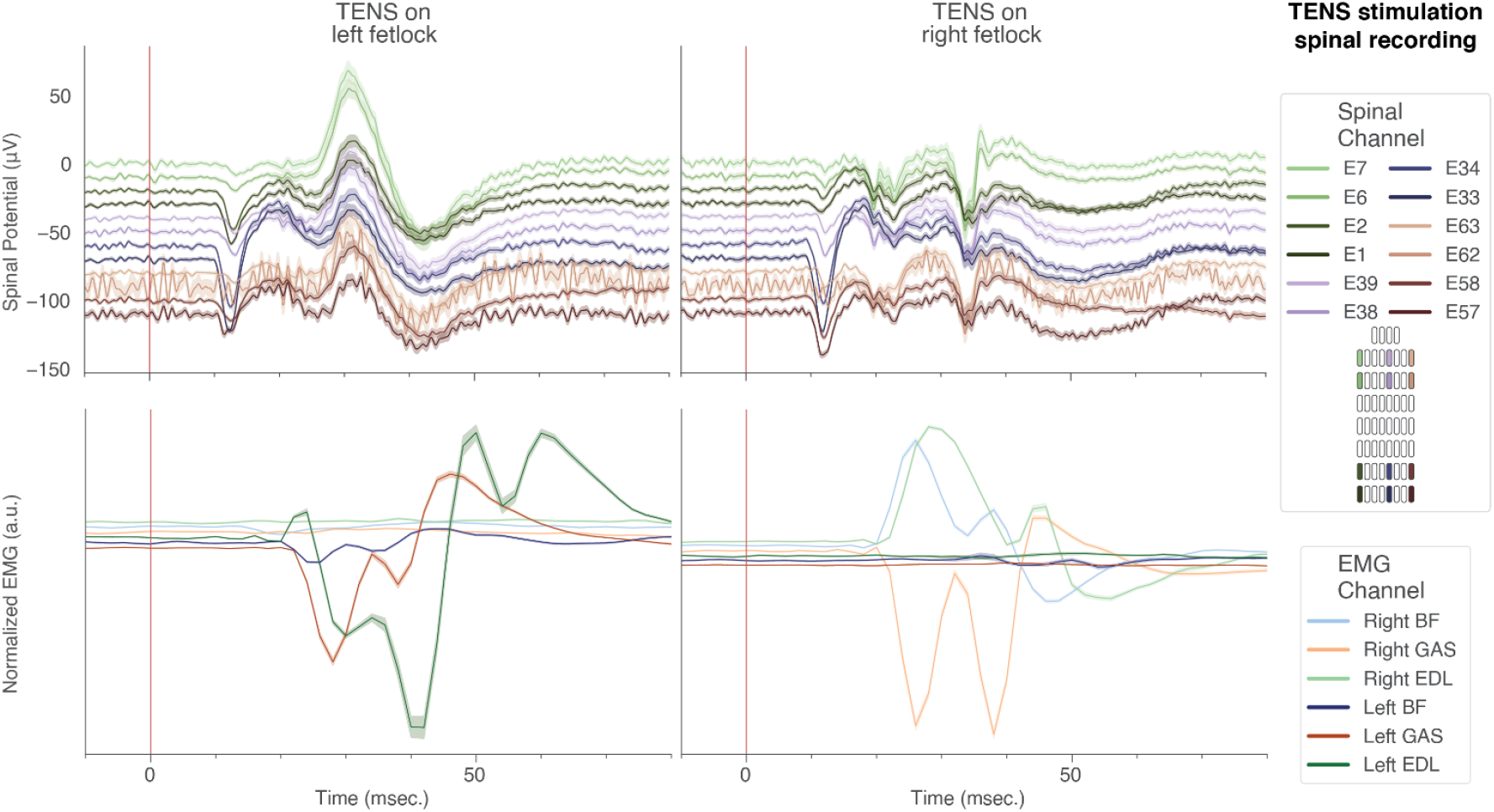
The MBI can resolve the spatiotemporal patterns of peripherally evoked spinal field potentials: mean and SEM (averaged across n = 50 pulses) of spinal ECAPs (top) and EMG (bottom) aligned to the onset of single TENS pulses (25V amplitude) indicated by a red line at t = 0 seconds.

### Stimulation response mapping

Finally, we used the MBI to record spinal ECAPs following electrical stimulation on the same electrode array used for the recording. Here, as well, EMG was recorded simultaneously to contextualize the ECAPs, especially with respect to latency after stimulation. Due to the presence of significant stimulation artifacts in such close proximity to the stimulating electrode, we employed a bipolar re-referencing scheme to the recordings, to better capture local effects of the stimulation on spinal potentials across the epidural surface of the cord. While the re-referencing process does not completely mitigate the slow return to baseline of the signal following the stimulation artifact, it reveals low-latency peaks in the spinal potential (fig. 7a,b) that are of putative neural origin [33], [39], due to their difference in latency.

**Figure 7.**
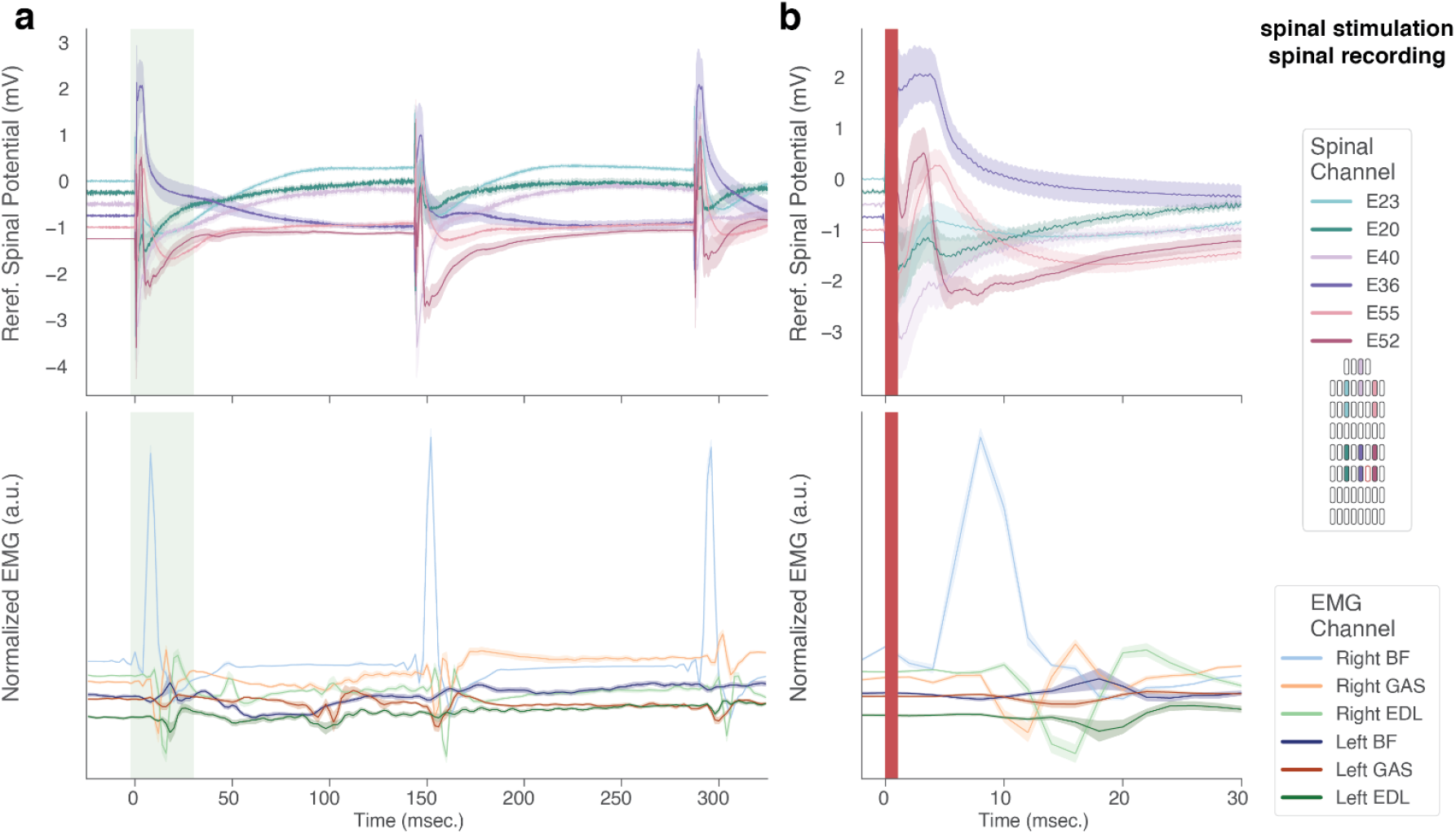
The MBI can resolve the spatio-temporal patterns of SCS-evoked spinal field potentials. (a) Mean and SEM (averaged across n=15 trials) of bipolar re-referenced spinal ECAPs (top) and EMG (bottom) aligned to the onset of the first stimulation pulse in a 300 msec train (8 Hz inter-pulse interval, 2.1 mA amplitude). (b) Detail corresponding to the time interval in panel e shaded in green. The stimulation artifact is masked by the red shaded region. Traces are vertically offset for ease of visualization.

## Discussion

Implantable devices capable of high-resolution bidirectional interfacing with the nervous system have the potential to revolutionize the treatment of neurological impairments and advance our understanding of the nervous system. Consequently, numerous research groups and private companies around the world have demonstrated great progress towards addressing the challenges associated with the development of such devices (table 1). Within that landscape, the MBI has five primary benefits: 1) recording resolution, 2) stimulation resolution, 3) capacity for closed-loop feedback by the implanted FPGA, 4) a high data rate wireless radio transmitting through the skin out of the body, and 5) a slim implant with a small footprint and modular extension leads adaptable to numerous sensing and stimulating leads. Here, we demonstrate the design of the MBI, verification of benchtop performance, and validation within a clinically relevant in-vivo model of spinal cord stimulation.

**Table 1.** MBI technical characteristics in relation to comparable implantable neural interfaces. * customizable from manufacturer ** streams action potential timestamps

| Device Name | Implant dimensions (mm) | Rec. BW (Hz) |  | chan. count |  | Effective data rate (Mbps) | ADC res. (bits) | sample rate (kHz) | On-board processing | Modular electrode | Ref. |
| --- | --- | --- | --- | --- | --- | --- | --- | --- | --- | --- | --- |
|  |  | low cutoff | high cutoff | rec. | stim. |  |  |  |  |  |  |
| MBI | 98.2 x 48 x 4.6 | 0.1 - 500 | 100 - 20k | 64 | 16 | 96.5 | 16 | 37 | Yes, FPGA | Yes | This work |
| SBNC | 56 x 42 x 9 | 0.1 | 7.8k | 100 | 0 | 24 | 12 | 20 | No | No | [13] |
| WiLD G1 | 51 x 37 x 5.5 | 0.1 - 500 | 100 - 20k | 128 | 0 | 61 | 16 | 30 | No | No | [62] |
| Link-32 | 31 x 23 x 6 | 0.1 | 500 | 32 | 0 | 0.768 | 12 | 2 | No | No* | [63] |
| Athena | n.r. | 0.3 | 7.5k | 32 | 32 | 11.52 | 12 | 2 - 10 | No | Yes | [64] |
| Neuralink | R 11.5 H: 8 | 5 or 300 | 10k | 1024 | 64 | 0.164** | 10 | 20 | Yes, spike detection | No | [4], [65] |
| Cortec Brain Interchange | 65 x 67 x 7 | 0.1 | 450 | 32 | 32 | 0.512 | 16 | 1 | No | No | [66] |
| WIMAGINE | R: 25 H: 12.54 | 0.5 | 300 | 64 | 0 | 0.45 | 12 | 1 | No | No | [20] |
| BISC | 12 x 12 x 0.05 | 4-55 or 1-22 | 15.1k | 256 or 1024 | 2 | 108 | 10 | 34 or 8.5 | No | No | [67] |
| RNS | 60 x 27.5 x 7.5 | 45759 | 30-120 | 4 | 4 | 0.01 | 10 | 0.25 | Yes, seizure detection | Yes | [68] |
| Summit RC+S | 57 x 47 x 6 | 0.5 - 8 | 100-2k | 4 | 16 | 0.195 | 15 | 1 | Yes, linear discriminant | Yes | [69] |
| Percept PC | 68 x 51 x 12 | 0.5 - 8 | 100-2k | 6 | 16 | n.r. | 15 | 0.25 | Yes, linear discriminant | Yes | [70] |

### The case for high-resolution closed-loop devices

There is a growing need for neural interfaces that offer a higher recording and stimulation channel count than current clinical devices. Traditional neuromodulation platforms for both spinal cord and deep brain targets rely on continuous injection of current at preset parameters [40], [41]. As these fields have advanced, closed-loop neuromodulation has gained traction, especially interventions where an electrophysiological signal is used as the feedback signal. There are numerous advantages to such closed-loop therapies.

For example, as patients progress through activities of daily living, the neural pathways engaged by the device can change. In the case of spinal cord stimulation, the physical distance between the stimulating electrode and cord can vary as a result of bodily movements. Newer spinal cord stimulation systems, such as the Saluda Evoke or Medtronic Inceptiv can account for this variance by continually sampling the size or shape of stimulation-evoked neural activity and tuning the stimulation intensity up or down to achieve a constant therapeutic effect [41]. The excitability of the neural tissue itself is also very dynamic, as a side effect of the complex interactions between its constituent neural circuits. During movement, as spinal reflex arcs modulate the state of different motor pools, identical pulses of electrical current can cause contractions of entirely different muscles [42]. This fact is critically important in the development of SCS for movement restoration after a spinal cord injury, where precisely timed pulses can enhance any residual descending motor commands [43].

Several challenges from the related field of deep brain stimulation reinforce the motivation for closed loop therapies. Many patients experience uncomfortable side effects from excessive stimulation amplitudes. Responsive algorithms might reduce the severity of these side effects, or at least continually adjust the stimulation based on the patient’s evolving needs. For example, DBS for movement disorders can often cause difficulties speaking. Closed loop DBS has been shown to reduce these effects, presumably by titrating the stimulation to a level that is still clinically effective while not causing off-target current spread [44]. As DBS has progressed to psychiatric applications, there has been much interest in both the clinical and scientific communities to identify plausible biomarkers of disease state, similar to beta oscillation power for movement disorders, that could serve as the feedback signal for closed loop algorithms [45], [46]. This search has been hampered by the relatively low sampling frequency and channel count of clinical-grade devices, when compared to state-of-the-art electrophysiological recording and stimulation systems, as one might find in a laboratory setting (see Table 1).

To summarize, converging evidence from several areas of neuromodulation suggests that increasing technical aspects of neural implants such as read/write channel count and temporal resolution can support the development of more effective therapies. The ratio of desired stimulation to recording channels is likely specific to each application. The MBI offers sixty-four recording channels and sixteen stimulation channels, which we believe to be an effective compromise that could be fine-tuned for specific applications. Note: The number of recording channels can be increased significantly by assembly of a lead with data communication capability (projected max channels of single unit recordings: 201 at 16-bit depth and 30kHz sampling). A higher maximum number of recording channels is provided than stimulation channels as many potential applications for closed-loop algorithms value recording more than stimulation coverage. Sufficient stimulation channels are certainly necessary for effective neuromodulation, particularly in cases where there are many potential off-target neural structures. In these cases, current steering can be used to limit the spread of activation only to the target area [47], [48]. However, several techniques can be employed to enhance the effectiveness of relatively few current sources.

Similar to recording channels, stimulation outputs can be used in conjunction with spatially multiplexed access to numerous stimulation sites to control the spread of stimulation current. Furthermore, sampling recording channels at rates of 30 kHz or faster provides access to signals that are otherwise unavailable such as action potentials. In contrast, due to the relatively slow timescales associated with the capacitance of neural membranes, on the order of hundreds of microseconds [49], [50], there is likely little benefit to delivering stimulation at faster rates. Instead, time-domain multiplexed delivery of current, where a single source sinks current into multiple sites very quickly, can mimic the presence of several independent current sources [51]. Ultimately, adding current sources to a device is subject to diminishing returns in terms of stimulation efficacy. In contrast, increasing the number of recording channels on a system can greatly improve the informational bandwidth of recorded signals, not only by increasing the Nyquist rate but also because new signal processing techniques become available with more channels, such as local re-referencing to remove noise [52] or geometric triangulation of signal sources [53].

### The benefits of a low-profile and modular system design

Here, we are presenting a slim implant with a small footprint that can be implanted on the cranium and throughout the body, efficiently using the limited available space for the total channel count and the supporting hardware for recording and stimulation. Development of a system that can be placed throughout the body presents challenges, as cranial mounted devices can potentially suppress cardiac artifacts compared to devices that are traditionally implanted in the torso [54]. Here, we demonstrate that the MBI system can obtain high-resolution *in vivo* recordings following chronic implantation in the torso. Furthermore, we demonstrate that the MBI can connect to third party devices, such as a high-resolution spinal cord stimulation electrode array. In the SCS field, electrode leads and connections are often used interchangeably across manufacturers due to similarity in size and pitch. However, there has been a concerted effort in the clinical neuromodulation field to follow the example of the cardiac pacemaker industry to establish standardized implanted connectors to minimize risk of orphan devices and facilitate interchanging components when appropriate. Therefore, development of a device-agnostic neural interface system such as the MBI will be necessary to effectively interrogate the nervous system across a range of devices.

Future versions of the proximal implant will be hermetically sealed for long-term use without an increase in thickness. The inductive power coil system of the MBI system indefinitely extends the lifespan of the implant beyond available battery power. One drawback of the power coil system is that magnets are required to locate the worn unit relative to the implant on skin. These magnets are not MRI-compatible, limiting current and future experiments.

### The benefits of UWB for data transmission

Over the past few decades, implanted medical devices have seen a dramatic increase in the amount of data that is being recorded and, therefore, needs to be transmitted to a receiver that lies outside the body. Traditionally, wireless data connectivity with implantable devices was achieved over the 402-405 MHz medical implant communication system (MICS) radio frequency (RF) band, following its standardization by the Federal Communications Commission (FCC) in 1999 [55]. However, MICS systems are limited to data rates on the order of 100 kbit/s, falling short of the capabilities of current research-grade neural recording systems which can simultaneously record high-bandwidth (20–30 kHz sample rate) high-resolution (10–16-bit ADC resolution) signals from hundreds of electrodes. Radio telemetry from an implanted device is challenging for several reasons, two notable examples being the absorption of radio signals by the body and interference from other electronic devices. The achievable data rate for any communication system is intrinsically proportional to the frequency of the carrier signal [56]. However, while radio waves in the 1-10 GHz range might be suitable from a bandwidth point of view, they are readily absorbed by the human body [57]. Furthermore, the radio spectrum is heavily regulated by governmental bodies such as the Federal Communications Commission (FCC) and the European Telecommunications Standard Institute (ETSI), which narrows down the range of frequency bands that could be used for telemetry from an implanted neural interface. Many existing standards for consumer electronics, such as Wi-Fi and Bluetooth, coexist in the industrial, scientific and medical (ISM) bands, which creates the risk of interference between devices and degraded functionality. To balance these multifaceted factors, we developed a dual-radio solution for the MBI, by combining a MICS band uplink with a UWB downlink. The older MICS standard is adequate for uplink operations that generally consist of updating operating parameters and, potentially, real-time control of electrical stimulation. Relatively few existing devices operate in the MICS band, therefore reducing the risk of interference.

Identifying a solution for high-bandwidth streaming of recorded data, on the other hand, remains an open question that is the subject of ongoing research [25], [58]. Here, we have opted for a UWB protocol that balances the requirements of high-data rate (96.5 Mbit/s) and low-power implanted device that has a low risk of interfering with existing radio devices [59], [60]. UWB technology is distinct from other RF communication systems in that the radiated energy is dispersed over a wide range of frequencies (3-10 GHz). As a result, UWB overlaps the 2.4 GHz ISM band used by consumer electronics, but falls under the regulated power cutoff defined by the FCC. Furthermore, the power in the overlapping band is a small fraction of the total, which greatly reduces the risk of interference. The major downside of UWB, resulting from its low spectral power density compared to, for example, Bluetooth, is a reduced range. However, in this application, the reduced range is irrelevant as the receiver is housed in the worn unit that must lie close to the implant in any case for wireless power transfer. Therefore, the close transmission range further reduces the risk of interference with other UWB-enabled devices.

UWB has several other benefits in this implementation beyond the high data rate. UWB’s wide distribution of power in the frequency domain corresponds to narrow pulses in the time domain, which has led to an alternative name for this technology, pulse radio. Interestingly, this fact underlies cybersecurity features that are not possible with comparable narrow band protocols. For example, UWB systems can precisely identify the distance between the transmitter and receiver by measuring the time-of-flight of the radio impulses.

Security protocols can make use of this fact and refuse connections to potentially malicious far-away devices [61]. As the cybersecurity of medical implants becomes the subject of increased scrutiny [62], support for enhanced security protocols becomes increasingly important. Finally, the functionality of precise spatial localization has led to increased adoption of UWB in recent years for commercial applications where tracking the location of assets is required. Consequently, miniaturized components (such as antennas) and small low-power integrated circuits for UWB telemetry have appeared on the market. Medical device development could benefit from this trend and utilize existing hardware to avoid the costly process of developing a new solution.

### Interpretation of in-vivo results

We could consistently communicate with the MBI system across the three months of implantation. However, the *in vivo* testing performed here was within a single sheep, and additional testing is required to verify the robustness of the surgical approach and system performance prior to formal clinical translation. The MBI was used with a worn unit in this application due to the ambulatory nature of experimentation. Future versions might offer optional configurations to reduce the worn load during *in vivo* evaluation. The worn unit reported here requires a cabled connection to power, future versions will offer a battery powered worn unit.

An ovine model was chosen for this demonstration as its spine is anatomically similar, albeit smaller than that of the average human [63]. Therefore, this model presented an opportunity to deploy clinical-sized devices in a laboratory setting and obtain proof-of-concept evidence for the device’s capabilities. Although the *in vivo* demonstrations reported here are limited to the epidural spinal cord, the small footprint and modular design of the MBI system could be utilized in micro or macro electronics in the central and peripheral nervous system in future applications. The MBI system was evaluated through three specific applications: spinal stimulation evoked motor responses, spinal stimulation evoked compound action potentials, and peripherally evoked compound action potentials. Evaluation of SCS electrode placement is commonly performed via stimulation of different contacts and mapping out areas of evoked motor activity or paresthesia [64]. Additionally, the use of ECAPs to titrate the delivery of SCS therapy has shown improved results compared to conventional SCS therapy, and achieved FDA approval in the US [65]. Furthermore, there has been recent interest in evaluating sensorimotor networks to neuromodulation by applying peripheral stimulation and recording centrally to understand the neurophysiological distribution of sensory and motor pools [66]. Therefore, the results shown here demonstrate that the MBI system has sufficient stimulation and recording resolution and modularity to detect known physiological signals from multimodal inputs.

## Conclusion

Here, we demonstrate the hardware design, benchtop validation, and in-vivo characterization of the Modular Bionic Interface (MBI). The MBI represents a significant advancement in the field of neural interfacing, combining high-resolution, bidirectional communication with modular design and wireless capabilities. Our benchtop evaluation confirms the performance of the MBI to component specifications and build requirements. Our in-vivo results demonstrate the capacity of the MBI to capture complex neural signals and deliver precise stimulation across experimental conditions. These features, paired with the scalability and adaptability of the MBI, highlight the system’s potential for broad applications in both research and clinical settings.

## Acknowledgments

We would like to thank Aaron Gregoire and the Center for Animal Resources and Education (CARE) staff at Brown University for their assistance with animal care. We would like to thank John Murphy for his engineering expertise in designing modifications to the sling apparatus.

## Authorship Statement

All authors contributed to the conception and design of the project. MM and IH developed the MBI system and designed and manufactured the hardware. NA, MM, and IH developed the MBI system firmware and its software platform. RD, JF, DB and ES performed the surgeries. RD, SP, JC, SS, ET, NA and MM acquired and preprocessed data. RD, SP, JC, ET, NA, MM, IH, and DB interpreted and analyzed data. RD drafted the initial manuscript. All authors edited the manuscript and reviewed the study results. Final approval was given by all authors.

## Data availability

The data that support the findings of this study are available at doi.org/10.17605/OSF.IO/9HXWE. Supporting code is available at https://github.com/rdarie/modular-bionic-interface-2026.

